# ProteoDisco: A flexible R approach to generate customized protein databases for extended search space of novel and variant proteins in proteogenomic studies

**DOI:** 10.1101/2021.09.17.460755

**Authors:** Wesley S. van de Geer, Job van Riet, Harmen J. G. van de Werken

## Abstract

**Summary:** We present an R-based open-source software termed ProteoDisco that allows for flexible incorporation of genomic variants, fusion-genes and (aberrant) transcriptomic variants from standardized formats into protein variant sequences. ProteoDisco allows for a flexible step-by-step workflow allowing for in-depth customization to suit a myriad of research approaches in the field of proteogenomics, on all organisms for which a reference genome and transcript annotations are available.

**Availability and Implementation:** ProteoDisco (R package version ≥ 0.99) is available from https://github.com/ErasmusMC-CCBC/ProteoDisco/.

**Supplementary information:** Supplementary table, figures and data files available.

## 1. Introduction

The rise and ease of current next-generation sequencing (NGS) techniques, coupled with reduced costs in both NGS and high-resolution mass-spectrometry, offers opportunity to incorporate sample-specific protein variants during proteomics experiments for increased accuracy and detection rates of, for instance, distinctive proteotypic peptides in bottom-up proteomics experiments. Expanding the repertoire of proteins and these proteotypic peptides can provide novel insights into disease-specific protein variants, their underlying molecular profiles and regulation, neoantigen prediction and expand our knowledge on the genetic variations encoded in proteomes.^1–5^ This is further fueled by the standardization and publication of proteomics resources which allows for the interrogation and combination of existing datasets.^6,7^

Rising global efforts in capturing the genetic sequences of diverse organisms, disease-related genotypes and their transcriptomes with subsequent proteome-resources warrants the implementation of a flexible yet intuitive toolset. This toolset should provide a bridge between genomic and transcriptomic variants and their incorporation within respective protein variants (proteogenomics) using industry-standard infrastructure, such as Bioconductor^8^, and allow for flexibility in facilitating the myriad experimental settings applied in research. Therefore, we designed and developed ProteoDisco, an open-source R software-package using existing Bioconductor class-infrastructures to allow for the accurate and flexible generation of variant protein sequences and their derived proteotypic peptides from the incorporation of sample-specific genomic and transcriptomic information. In addition, we present the results of ProteoDisco and two similar open-source tools which are frequently utilized within proteogenomics (customProDB^9^ and QUILTS^3^) with their performance in generating correct protein variants and respective proteotypic peptides from supplied genomic variants.

## 2. Approach

ProteoDisco incorporates genomic variants, splice-junctions (derived from transcriptomics) and fusion genes within provided reference genome sequences and transcript-annotations to generate their respective protein variant sequence(s). These sequences can be curated, altered and subsequently exported into a database in FASTA-format for use in downstream analysis. To limit the number of generated protein variants, ProteoDisco provides filtering options based on a minimal number of distinct proteotypic (identifiable) peptides. The global workflow of ProteoDisco is summarized in six steps as depicted within **Figure 1**. In addition, an extended overview of how (novel) splice-junctions and gene-fusion events are incorporated is shown in **Supplementary Figure 1**.

**Figure 1.**
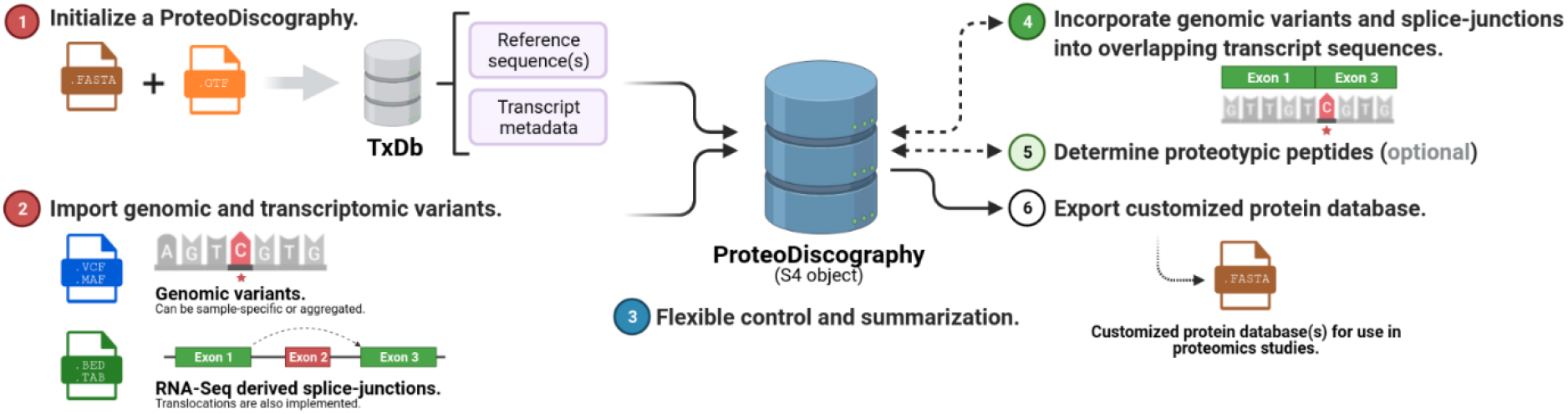
schematic overview of the ProteoDisco workflow. The global workflow of ProteoDisco can be categorized as six major steps. 1) Initialize a ProteoDiscography by utilizing custom references sequence(s) and gene-annotation(s) or using pre-existing TxDb objects. 2) Import (sample-specific) genomic variants, splice-junctions or manual sequences. Several sanity-checks are performed during importation, including the validation of matching reference nucleotide(s). 3) Dynamically view, extend, alter and customize imported records and derived sequences. 4) Incorporation of genomic variants and splice-junctions into overlapping transcript-annotations, translocations between chromosomes can also be processed. The incorporation can be performed in a sample-specific manner, exon or transcript-specific manner or in an aggregated manner. 5) Cleave derived protein variant sequences and determine proteotypic peptides, per protein, which are not present within the reference protein sequences (TxDb) or additional protein databases (e.g., UniProKB). 6) Export the derived protein variant sequences into an external protein-sequence database(s) using FASTA format.

To compare the accuracy of ProteoDisco against two common alternatives for proteogenomics studies (customProDB^9^ and QUILTS^3^), we utilized a manually-curated dataset and two large independent proteomics studies. The manually-curated dataset contained 28 genomic variants reported in COSMIC^10^ comprising multiple variant classes; synonymous and nonsynonymous single-nucleotide variants (SNVs), multi-nucleotide variants (MNVs) and in- and out-of-frame insertions/deletions (InDels). In addition, we utilized recently-published results from large-scale colon and breast cancer cohorts within the Clinical Proteomic Tumor Analysis Consortium (CPTAC) to illustrate the accuracy of ProteoDisco in generating identical proteotypic peptides as detected within these studies.^2,5^ This comparison revealed that ProteoDisco correctly generated proteotypic peptides from their respective genomic variants after thorough checking and yielded the highest number of expected and reconstructed proteotypic peptides within all three datasets (**Supplementary Figure 2**). In total, only four enigmatic genomic variants (of three fragments) from Mertins *et al*. could not be reconstructed to reproduce their proteotypic peptide(s).

## 3. Conclusion

In this article, we present ProteoDisco, a suitable, open-source and flexible suite for the generation of protein variant databases usable in downstream proteogenomic studies and capable of correctly incorporating a diverse range of genomic variants and transcriptomic splice-junctions. We report that ProteoDisco accurately produces protein variant sequences harboring previously-identified proteotypic fragments from their respective genomic variants. Further examples and use-cases can be found in the vignette of the ProteoDisco package.

## 4. Methods

### 4.1 Technical design of ProteoDisco

ProteoDisco was programmed within the R statistical language (v4.1.1) and built upon existing classes within the Bioconductor infrastructure (v3.13) to allow flexible inheritance and future extensions. Additional information on the usage and design of ProteoDisco can found in the extended methodology (**Supplementary File 1**).

### 4.2 Assessment of the correct integration of genomic variants into protein variants

We generated a custom validation-dataset containing established somatic variants (SNVs, MNVs and InDels; *n* = 28) and their respective protein variants as listed within COSMIC^10^ (v92; GRCh37; **Supplementary Table 1**). In addition, we utilized recent proteogenomics studies from the CPTAC cancer cohorts containing genomic variants and their respective *in silico* generated proteotypic peptides which had been measured and identified using high-throughput proteomics approaches.^2,5^ In the Wen et al. dataset^5^ (CPTAC - Colon Cancer), genomic variants (and their respective proteotypic peptides) were split into sample-specific VCF-files based on the data present within their published Suppl. Data S1 (sheet 1: ‘prospective_colon_label_free_in’). The Mertins et al. dataset^2^ (CPTAC - Breast Cancer) was aggregated into a single VCF-file based on the data present within their published Suppl. Table S5 (sheet 2: ‘Variants’).

Using these three datasets, we ran ProteoDisco (v0.99), customProDB (v1.30.1) and the web-interface of QUILTS (v3.0; as available from http://openslice.fenyolab.org/cgi-bin/pyquilts_cgi.pl; accessed 13-04-2021) to generate custom protein-variant databases using uniform UCSC/RefSeq^11^ (GRCh37) transcript-annotations and settings. The custom protein-variant databases were generated based on two approaches within ProteoDisco. The first approach incorporated each genomic variant independently and the second allowed for the simultaneous incorporation of all genomic variants per overlapping transcript-annotation, e.g., two variants on different coding exons would both be incorporated within the resulting variant protein-sequence. Incorporation of all possible combinations of mutant exons yields too many combinations and is therefore not included amongst the options.

The generated variant protein sequences and respective proteotypic peptides from each customized protein-variant database were compared against the proteotypic peptides as expected from COSMIC or as detected within the respective CPTAC-studies using all three tools (**Supplementary figure 2**). E.g., if ProteoDisco generated three distinct proteotypic peptides for a given genomic variant and one of those was identified within CPTAC (or COSMIC), it was counted as a concordant result.

### 4.3 Code availability

All source-code has been deposited within GitHub (https://github.com/ErasmusMC-CCBC/ProteoDisco) under the GPL-3 license and has also been made available within Bioconductor (**[currently under submission](https://github.com/Bioconductor/Contributions/issues/2206)**).

### 4.4 Data availability

The custom validation dataset (GRCh37) which has been used in the analysis as presented within this manuscript has been stored within ProteoDisco and is accessible at https://github.com/ErasmusMC-CCBC/ProteoDisco/main/inst/extdata. COSMIC (v92; accessed on 14-04-2021) was used to derive the validation dataset (GRCh37), the external validation datasets based on CPTAC (colon and breast cancer) were generated based on the supplementary data published by Wen et al.^5^ and Mertins et al.^2^.

## Author contributions

All authors had full access to all the data in the study and take responsibility for the integrity of the data and the accuracy of the data analysis.

*Study concept and design:* van Riet, van de Geer, van de Werken.

*Acquisition of data:* van Riet, van de Geer.

*Analysis and interpretation of data:* All authors.

*Drafting of the manuscript:* All authors.

*Critical revision of the manuscript for important intellectual content:* All authors.

*Statistical analysis:* All authors.

*Obtaining funding:* None.

*Administrative, technical, or material support:* All authors.

Supervision: van de Werken.

Other: None.

## Acknowledgements

We would like to thank S. Eldeb and W. M. Hoska for their preliminary work on the integration of mutations into transcript sequences. **Figure 1** and **Supplementary Figure 1** were created using BioRender.com.

## Supplementary Tables

**Supplementary Table 1 - Overview of comparisons**.

Extended overview of the comparisons of three variant datasets on ProteoDisco, customProDb and QUILTS.

## Supplementary Figures

**Supplementary Figure 1 - Overview of the procedure of generation mutant splice-isoforms based on translocations- and non-canonical splice-junctions**.

Schematic overview on the handling of splice-junctions (SJ) to generate splice-isoforms. Optionally, users can opt to only generate non-canonical splice-isoforms and fusion events, thereby ignoring canonical forms already present within the ProteoDiscography TxDb.

**Supplementary Figure 2 - The number of concordant proteotypic peptides for ProteoDisco, customProDb and QUILTS for our manually-curated test-set and two CPTAC-datasets (colon and breast cancer)**.

Venn-diagrams displaying the absolute number and relative percentage of identical proteotypic fragments after incorporation of genomic variants, per dataset and tool. We tested ProteoDisco (v0.99), customProDb (v1.30.1) and QUILTS (v3.0) using uniform annotations and settings.

a) Overlap of concordant results based on our validation dataset (COSMIC; GRCh37).

b) Overlap of concordant results based on the CPTAC colon cancer dataset (Wen et al.).

c) Overlap of concordant results based on the CPTAC breast cancer dataset (Mertins et al.).

## Supplementary Files

**Supplementary File 1 - Extended Materials and Methodology on the design of ProteoDisco**.

The extended materials and methodology detailing the technical design of ProteoDisco.

## Notes

### Competing Interest Statement

The authors have declared no competing interest.

