## Supplementary figures and images for "ProteoDisco: A flexible R approach to generate customized protein databases for extended search space of novel and variant proteins in proteogenomic studies"

### Supplementary Figure 1

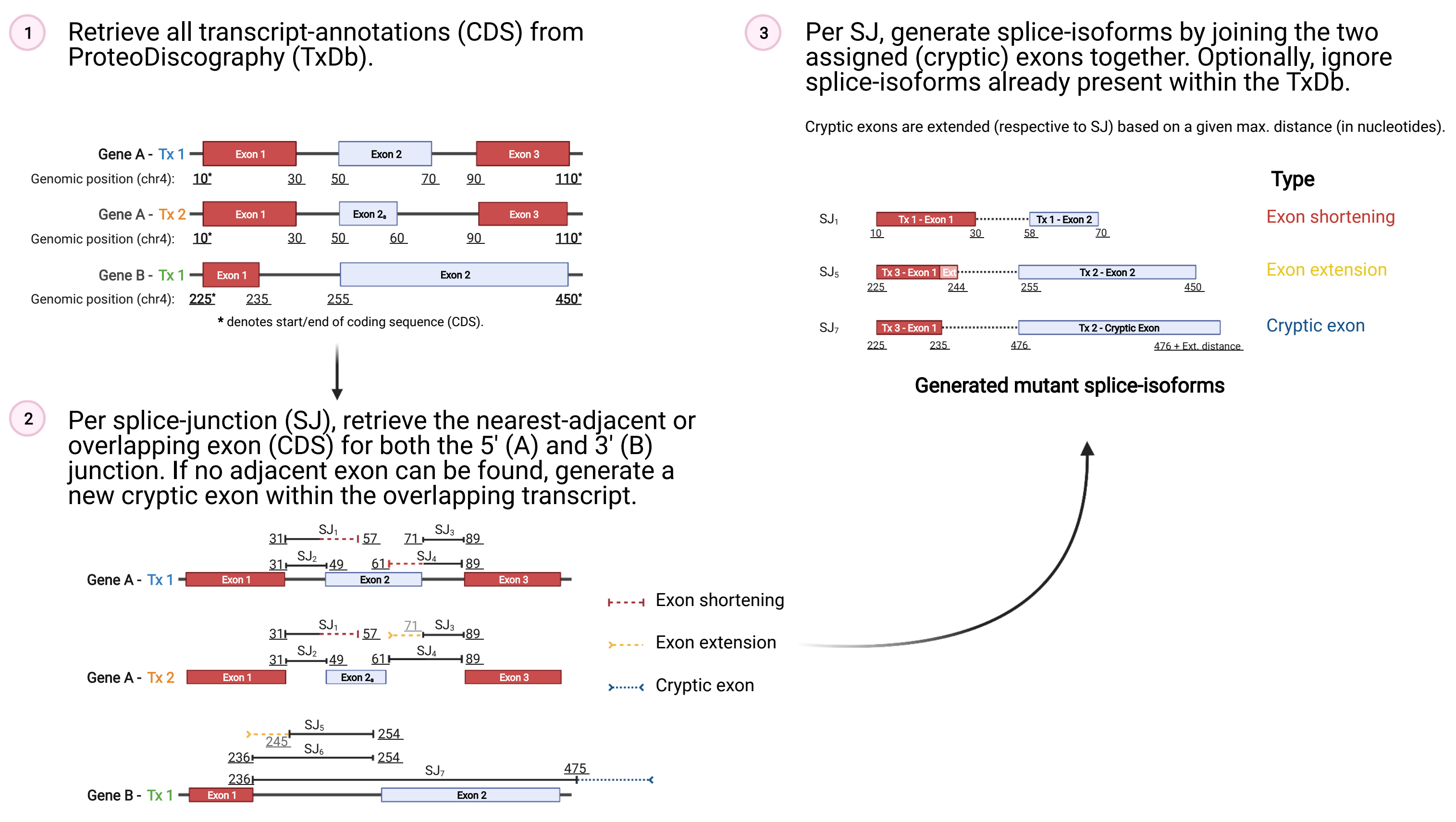

### Supplementary Figure 2

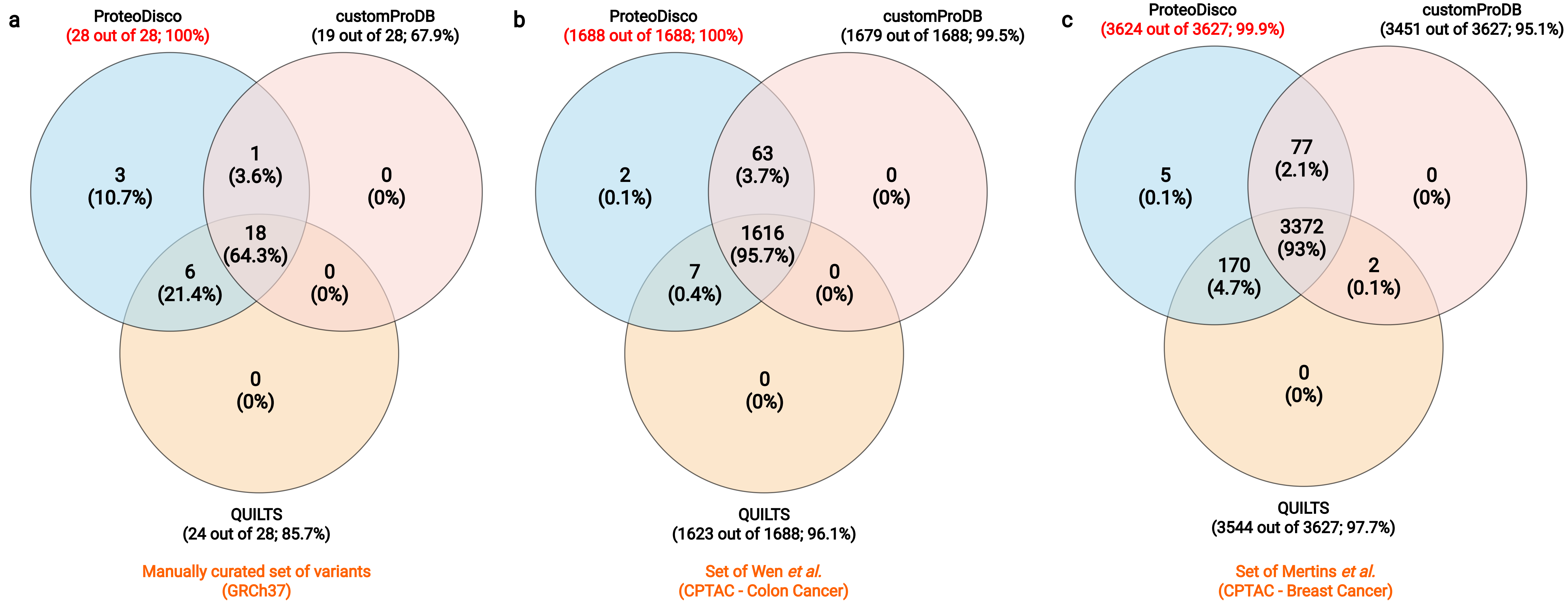
