## Supplementary Methods for "ProteoDisco: A flexible R approach to generate customized protein databases for extended search space of novel and variant proteins in proteogenomic studies"

### **Supplementary File - Extended Materials and Methodology on the design of ProteoDisco.**

The major workflow of ProteoDisco can be divided into six steps;

1. Generation of the ProteoDiscography containing the reference genome sequences and transcript annotations of choice, as detailed below. This ProteoDiscography will also house the imported genomic and transcriptomic input and subsequent in silico generated protein variants and related information.
2. Import of genomic variants (either VCF and MAF files or VRanges objects12) or splice-junctions from transcriptomics such as .BED output from TopHat13, SJ.out.tab output from STAR14 or manual entries following a simple format to for instance denote translocations and/or fusion-gene events (e.g., TMPRSS2-ERG). ProteoDisco is capable of handling SNVs, InDels, MNV variants of both non-synonymous as synonymous variants.
3. Integration of genomic variants and splice-junctions into their respective transcripts and coding sequences (CDS) to generate in silico transcript variants, as detailed below.
4. Translation of in silico generated transcript variants into their respective protein variants, the genetic code used for translation can be altered to allow for divergent translation tables for non-standard orgasms.
5. Determine the number of proteotypic peptides per transcript variant, this can be determined based against the given reference database (as given to the ProteoDiscography) or be extended with additional protein-sequence databases. In addition, ProteoDisco can also check for proteotypic peptides compared to the other generated protein variants.
6. Export of the generated protein variants into a distinct FASTA database for use in downstream proteomics analysis to extend the (sample-specific or cohort-wide) search-space.

#### **1. Generation and design of the ProteoDiscography; the internal data-structure.**

All reference genome sequences (BSGenome objects), transcript annotations (TxDb objects) and generated results (BioStrings, tibbles and DataFrames) throughout ProteoDisco are housed within a custom (S4-class) termed ProteoDiscography. The reference database and transcript annotations for the ProteoDiscography can be generated in two ways; using pre-generated BSGenome (reference sequences) and TxDb (transcript annotations) objects, for instance available from BioConductor, or by supplying the reference genome sequences and transcript annotations (FASTA and GTF/GFF file, respectively) which in turn generates these objects. In addition, the genetic code can be specified (as detailed by Biostrings) to also allow for non-standard translation tables.

#### **2. Import genomic variants and splice-junctions within the ProteoDiscography.**

After initialization of a ProteoDiscography, genomic variants and splice-junctions can be imported.

Genomic variants (or somatic mutations) can be imported from .VCF or .MAF files or VRanges objects containing the genomic positions, strand and reference/variant alleles. By default, all given reference anchors (genomic position(s) and reference allele) of the genomic variants are checked against the provided reference genome and nucleotide at the respective position(s) to prevent inconsistencies. If non-matching reference anchors are detected, ProteoDisco will either halt the import-process and whilst displaying the erroneous records or, by setting ignoreNonMatch = TRUE, it will report and remove these non-matching records and continue with the remainder.

Splice-junctions can be imported from standard .BED (e.g., TopHat) and.SJ.out.tab (e.g., STAR) files or manually supplied using a simple format. Each of these formats should detail the genomic position (and optionally, strand information) of the donor and acceptor junction-sites for each splice-junction (junctionA and junctionB, respectively). Manual input can be supplied using the following format:

- junctionA: Genomic coordinates of the 5'-junction (i.e., the position of the first intronic base).
  - Format: chr:start:strand, i.e.: chr1:100:+
- junctionB: Genomic coordinates of the 3'-junction (i.e., the position of the last intronic base).
  - Format: chr:start:strand, i.e.: chr1:150:+
- sample: Sample-identifier. (optional)
- identifier: Identifier for the splice-junction, this identifier will be used to denote the splice-junction in downstream analysis. (optional)

This manual-input can be also be used to supply splice-junctions from translocation events such as BCR-ABL which result in a protein variant containing exonic sequences from two chromosomes.

In addition, users can also supply pre-determined full-length transcript sequences into the ProteoDiscography. These manually-supplied transcript sequences can then also be used to determine proteotypic peptides compared to the reference database and/or protein variants.

ProteoDisco houses functions to detect duplicate samples and overwrite these (if required) or append new genomic variants and/or splice-junctions to existing samples (based on sample names). In addition, it can also be toggled to remove all pre-existing samples within the ProteoDiscography prior to importation of new input.

#### **3. Incorporation of genomic variants and splice-junctions within the coding sequence of overlapping transcripts.**

ProteoDisco facilitates options to incorporate all supplied genomic variants (incl. synonymous variants) for all samples simultaneously or to perform this on a per-sample basis (aggregateSamples). Similarly flexible, users can choose between incorporating all mutations (per-sample or all samples aggregated) within the same transcript (e.g., a single RNA transcript containing 5 mutations; aggregateWithinTranscript = TRUE) or to generate separate transcripts, each harboring only a single mutation (e.g., 5 transcripts for 5 mutations; aggregateWithinTranscript = FALSE). Finally, users have similar functionality at exon-level (aggregateWithinExon).

Based on the parameters set by the user, genomic variants are overlapped with the coding sequences (CDS) of each transcript within the supplied TxDb. Per variant, all overlapping CDS (from one or multiple transcripts) will be altered by incorporating the overlapping genomic variant(s) at the correct coding position. The reference anchor (reference allele) will be checked if this conforms to the nucleotide at the coding position, taking in mind the orientation of the CDS. Genomic variants (e.g., InDels) overlapping the intron-exon or exon-intron boundary of a CDS will be split and only the CDS-overlapping portion will be incorporated.

After all genomic variants have been incorporated within their overlapping CDS in the transcript(s), the transcript sequence is generated by stitching all CDS of the transcript from 5’ to 3’ together. Based on the parameters set by the user, this will either results in a single transcript variant containing all mutant CDS or multiple transcripts with distinct mutant CDS. In addition, the 3’ untranslated region (3’ UTR) is also added to the mutant transcript sequence to capture additional coding nucleotides after a possible loss of the canonical stop-codon.

Splice-junctions are handled by determining the nearest adjacent (5’ and 3’) or overlapping CDS sequences. If a splice-junction overlaps with an existing CDS, that CDS will be used as assigned CDS and be altered to start (if 5’; junctionA) or end (3’; junctionB) at the genomic position of the respective junction, resulting in a shortened CDS. If the junctions do not overlap with an existing CDS, it will be assigned to the nearest adjacent CDS, taking in mind the orientation and strand of the splice-junction and CDS. This will either retrieve a canonical CDS directly flanking the splice-junctions or assign a CDS further away. The nucleotides spanning the splice-junction to the assigned CDS will then be added to assigned CDS and thereby effectively extending the CDS. If the splice-junction is further away than a max. distance (as set by the user; default 250 nt), a cryptic exon (of a size set by the user; default = 99 nt) will be generated and incorporated within all overlapping transcripts. As we cannot discern frame-status for cryptic exons, a three or six-frame translation (if splice-junction has no strand information) will be performed.

The generated splice-junction-derived transcript can also span two distinct genes; e.g., if one junction is most adjacent to gene X and the second junction is most adjacent to gene Y. These ‘fusion’-genes are then generated by stitching (taking the strand into account) of all upstream CDS of gene X (ending at the assigned 5’ CDS) with all downstream CDS of gene Y (starting at the assigned 3’ CDS); extensions and/or shortenings are also incorporated in these situations.

Post-incorporation, all generated transcripts (genomic variants, splice-junctions and/or manual sequences) can be curated and altered using the setMutantTranscripts function.

After the generation of transcript variants (or manual alteration thereof), all mutant transcript sequences are translated into their respective protein sequences and cleaved at the earliest stop-codon. If the canonical stop-codon is lost, it will continue translating into the 3’ UTR until the next-earliest stop-codon or stop at the end of the 3’ UTR.

Generated splice-junctions transcripts without known translation frame(s) will generate a three-frame (if orientations are known and concordant) or a six-frame (if strand orientations is unknown or disconcordant) translation of the transcript sequence(s).

#### **4. Filtering for protein variants based on proteotypic peptides.**

To reduce number of potential protein variants, ProteoDisco provides an optional filtering procedure to retain protein variants containing a min. number of proteotypic peptides not seen in the supplied TxDb (and additional) protein databases and thereby identifiable in subsequent MS/MS analysis.

Conceptually, we cleave the protein variants with the same protease as would be used in the respective MS/MS experiment (e.g., Trypsin) and compare the resulted cleaved peptides against the input TxDb (and additional databases) cleaved in the same manner (allowing user-set missed cleavages) and, subsequently, determine the number of distinct cleaved fragments not detected in the reference protein database(s). In addition, it can also be toggled to check for uniqueness against all other generated protein variants within the ProteoDiscography. This extend the ProteoDiscography with the number of proteotypic peptides per protein variants which can be used to filter protein variants prior to exporting the protein sequences to a FASTA file (step 5).

#### **5. Export protein variants into a customized protein database (FASTA).**

Generated protein variants can be exported into an external (FASTA) database. As mentioned, users can subset exported proteins based on the minimum number of proteotypic peptides. Users can output all generated protein variants into the same aggregated file or generate distinct files containing sample-specific protein variants.

The FASTA headers for each protein sequence contain identifiers and information on the incorporated variant(s) or splice-junctions which can be easily related back to the ProteoDiscography.
